# Amino acid oxidation constrained protein sequences during evolution: an analysis of a model bacterium

**DOI:** 10.1101/2020.06.24.169870

**Authors:** Enrique González-Tortuero, Alexandro Rodriguez-Rojas

## Abstract

Within the cell, there are compartments and local conditions that may constrain amino acid sequence. Life emerged in an anoxic world but releasing a photosynthesis by-product, the molecular oxygen, forced adaptive changes to counteract its toxicity. Several mechanisms evolved to balance the intracellular redox state and maintain a reductive environment more compatible with many essential biological functions. Here, we statistically interrogate the aminoacidic composition of *E. coli* proteins and investigate how the proneness or susceptibility to oxidation of amino acid biased their sequences. By sorting the proteins in five different compartments: cytoplasm, internal membrane, periplasm, outer membrane and extracellular; we found that various oxidative lesions constrain protein composition, that there is a dependency of the cellular compartments and there is an impact on the evenness distribution or frequency. This analysis is necessary from an evolutionary point of view to reflect how the oxidative atmosphere could restrict protein composition and probably imposed a trend in the codon bias.

## Introduction

There is still much mystery in the origin of life and pristine molecular evolution. During molecular evolution, amino acid sequences were selected from their functionalities^1^, stability^2,3^, energy economy^4,5^ and the ability to create secondary structures^6^. This process was a dynamic mutationselection game that gave particular combinations of amino acidic sequences^7^. Proteins are the ultimate product of the genetic flow information. In the last instance, protein functions are determined by the amino acid sequence. However, these functions must be harmonised with the biological complexity, heavily influenced by environmental conditions^8^. Nevertheless, live evolved in an anaerobic and reductive atmosphere, and oxygen generated by photosynthesis changed life forever and forced the microbes to alter the physiology to cope with oxidative conditions^9,10^. Oxygen and its reactive species (ROS) can damage all cellular components such as lipids, nucleic acids and proteins^11,12^.

Inside the cell, the cytoplasm is kept under reductive conditions due to the evolved systems to control ROS^13^. ROS damages all cellular components, including proteins, and those oxidative modifications compromise protein biological activities^12^. The high frequency of protein oxidation forces the cell to perform the turnover that speeds up in the presence of an excess of oxidative agents^13,14^. As proteome quality control, several proteolytic and chaperone systems cooperate in ridding off the non-functional and structurally altered proteins^15^. Nevertheless, secreted proteins, which play essential functions for bacteria, escape such quality control, and they probably encounter more adverse conditions, including more oxidative environments.

By contrast, the oxidation of cytoplasmic and membrane-associated proteins is tightly associated with ageing. Indeed, until recent years it was thought that microbes were immortal or refractory to ageing processes due to the binary division that theoretically generates two identical cells^16,17^. After division, the splitting of bacterial proteins is asymmetric, and such asymmetry correlated with the accumulation of oxidised proteins and, in the last instance of bacterial ageing, death^18–20^. For example, carbonylation leads to protein aggregation and intracellular precipitation, which is harmful to the cell, associated with senescence and increases the probability viability loss^17^.

There is a redox potential gradient going from intracellular compartments to extracellular medium in the direction of from reductive to an oxidative state that may occur in an oxygen-rich scenario. This compartmentalised gradient is less complex in Gram-positive and by cytoplasm, cell envelopes and extracellular space. In Gram-negative bacteria, there are two additional compartments due to the presence of the outer and inner membranes. Both membranes generate a periplasmic space. In a previous study, optimisation of extracellular proteins to reduce their energetic synthetic cost was considered taking into cellular compartments^5^. Here, we hypothesise that redox states of the micro-environment constrain the amino acid composition of proteins across cellular compartments in prokaryotic cells. Thus, we stressed this possibility by analysing the frequencies of amino acids of every protein in the whole proteome of the model bacteria *Escherichia coli*, normalised by the expression level, and their correction with oxidation events.

## Results and Discussion

We first compiled all amino acid sequences belonging to proteins classified by cellular compartment: cytoplasm, inner membrane, periplasm, outer membrane and extracellular environment (Table S1). For such purpose, we took into account for our analyses those amino acids that are more prone to oxidative lesions using as a model the well-annotated *Escherichia coli* genome K12 MG1655. Unfortunately, these type of analyses would be compromised on other microorganisms which information on protein location is not well known and not given a function^21^. These comprise methionine sulfoxidation and disulfide formation, histidine, tyrosine and tryptophan oxidation, peroxidation, adduct formation, metal-catalysed oxidation, and carbonylation^22^. The carbonylation is probably the most frequent one in cells^23^. There is an heterogeneity in number of proteins per cellular compartment in *E. coli* K-12 MG1655 (Cytoplasm: 2689; Inner Membrane: 941; Periplasm: 349; Outer Membrane: 146; Extracellular: 16; Fig. 1). The imbalance of protein distribution numbers is a potential source of bias across the analyses.

**Figure 1.**
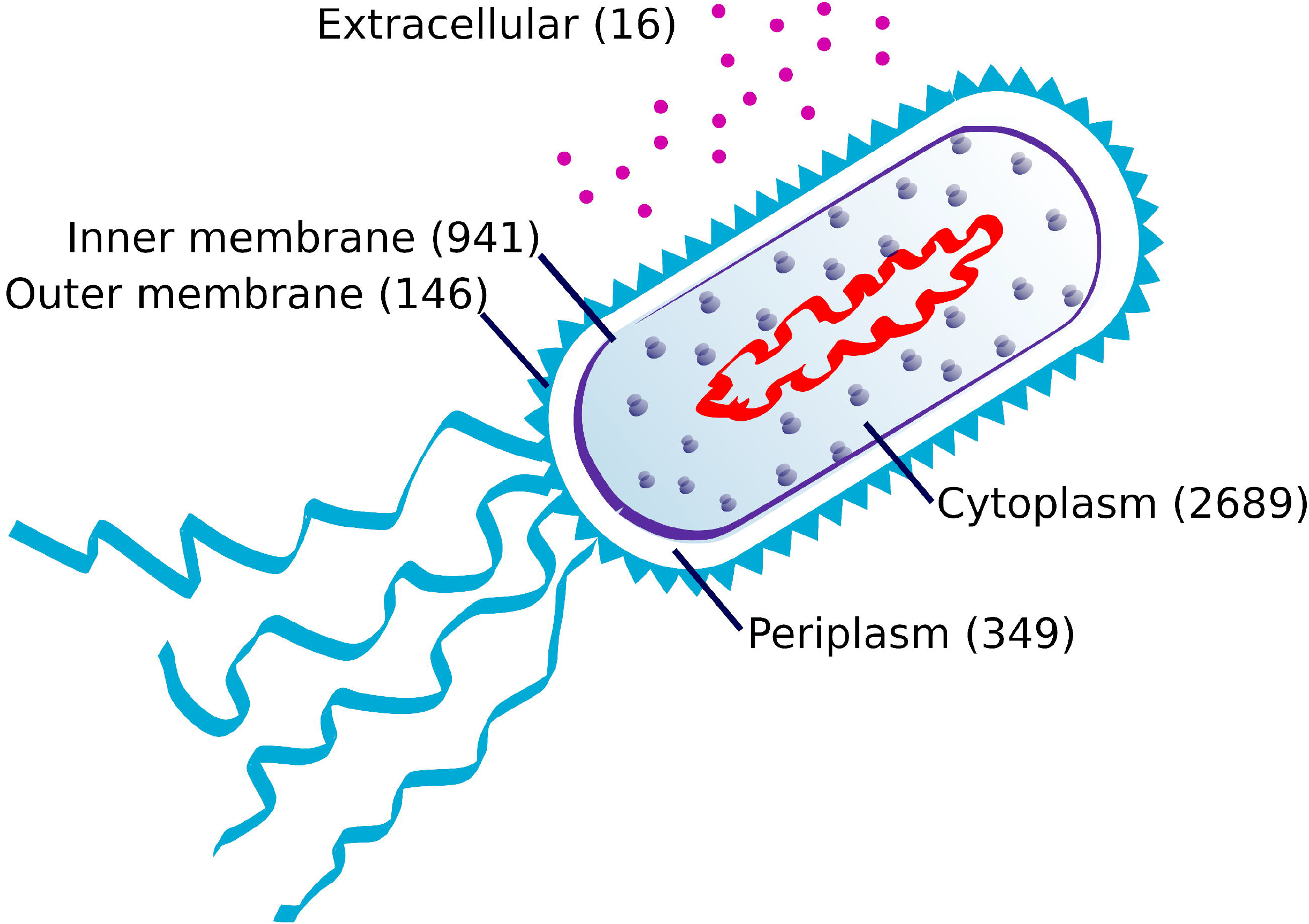
Graphic representation of the total number of *E. coli* proteins segregated by compartments (cytoplasm, inner membrane, periplasm, outer membrane, extracellular media). Numbers between parenthesis indicates the number of proteins in each compartment. We assigned to the extracellular media only secretable proteins, for example flagella proteins are classified as inner membrane, periplasmic or outer membrane proteins depending on their location.

A neutral analysis of protein amino acid composition reveals that, in general, there are significant differences for all amino acids (GLM: *P* = 0.0283). Only alanine, aspartic acid, isoleucine, lysine, leucine, methionine, proline and glutamine did not show any preference for any cellular compartment (Table S2). Similar results were found in the comparison of 38 proteomes among the tree of life, whereas low reactive amino acids—glycine, alanine, isoleucine and valine—were predominant in proteins with a prolonged half-life^24^. Thus, we focus our attention on susceptible amino acids residues. In all cases, there is a decrease in the content of vulnerable amino acids across subcellular compartments (Fig. 2). Based on our hypothesis that amino acid susceptibility to oxidation might constrain protein sequences, the distribution of the frequency of amino acids for all five compartments was analysed. We understand that this correlation does not imply causation but allow us to hypothesise about disparities on the amino acids sequences of proteins across compartments. Due to the nature of the data, no experimental approaches are available to test this hypothesis as happened in previous studies^4,5,24^.

**Figure 2.**
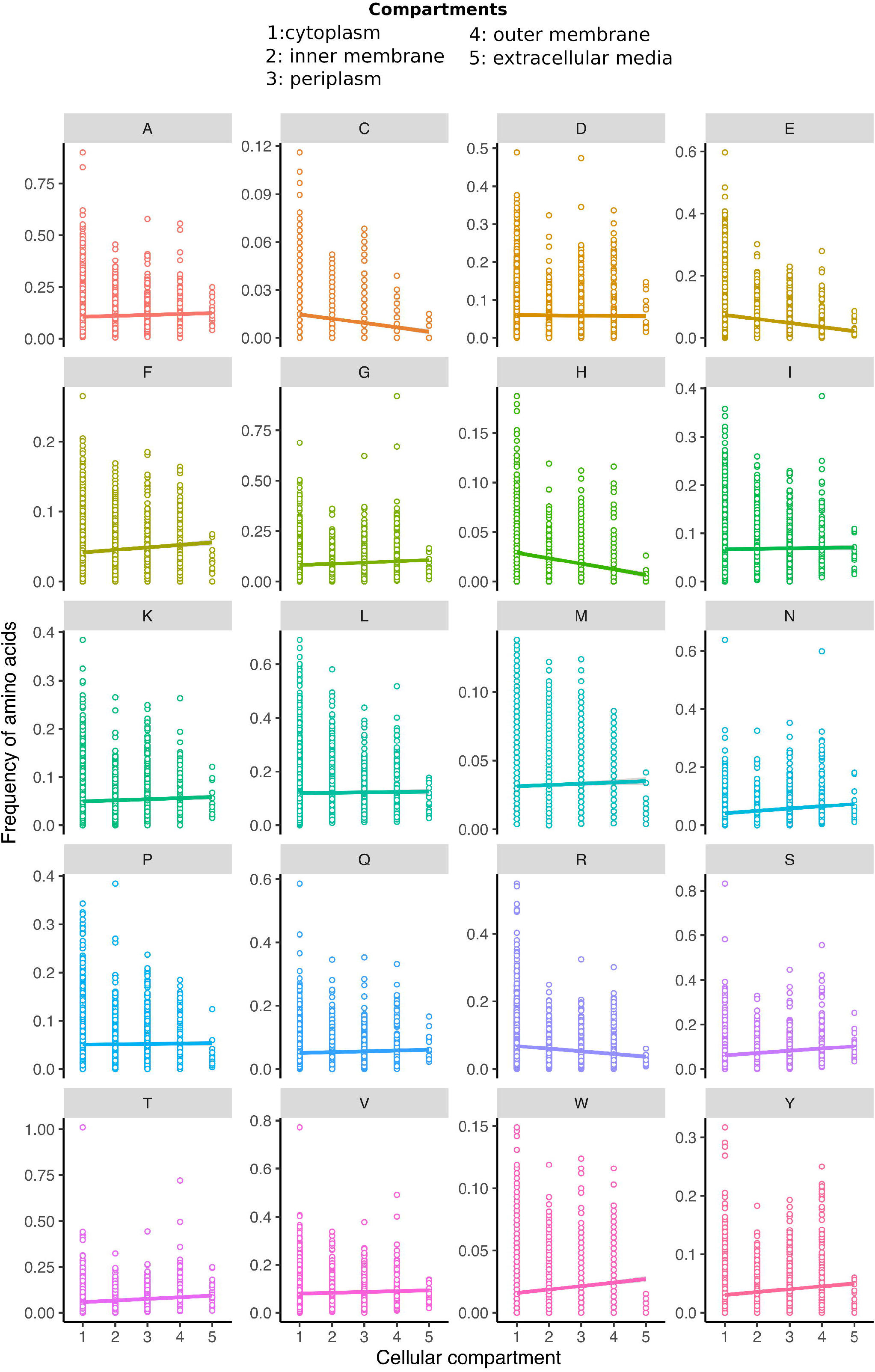
Distribution of the frequency of every amino acid within proteins for all subcellular compartments (Cyt: cytoplasm, IM: inner membrane, Per: periplasm, OM: outer membrane, Ext: extracellular media). Note that the scale for each amino acid occurrence frequency was adjusted to the maximum values and is different for each case. The trend lines represent GLM model which is significant for all amino acids except for alanine, aspartic acid, leucine, isoleucine, lysine, methionine, proline and glutamine (to consult statistic values see Table S1).

### Aberrant disulfide bonds formation susceptibility

Cysteine helps to maintain the redox state in the cytoplasmic compartment^25^. When it comes to cysteine frequency categorised in compartments, there is a significant difference between cytoplasmic proteins and the rest of compartments—from the inner membrane to the extracellular space; Kruskal-Wallis Test: *P* < 2.200×10^-16^, Mann-Whitney *U* (Cytoplasm – Inner membrane): *P* = 9.764×10^-35^, Mann-Whitney *U* (Cytoplasm – Periplasm): *P* = 2.615×10^-10^, Mann-Whitney *U* (Cytoplasm – Outer membrane): *P* = 2.714×10^-5^, Mann-Whitney *U* (Cytoplasm – Extracellular): *P* = 1.342×10^-4^. In fact, there is a trend to the reduction in the content of cysteine going from intracellular to extracellular compartments—GLM: slope = −2.717×10^-3^; *P* < 2.200×10^-16^; Spearman correlation: *r_s_* = −0.205; *P* < 2.200×10^-16^ (Fig. 3A). Covalent linking of amino acids side chains within a polypeptide adds to the stability and function of several proteins, being these disulfide bridges the most common^26^. However, the possibility of aberrant disulfide bonds can cause mispairing of cysteines leading to misfolding, aggregation and irreversible oxidative damage^27^. In general, disulfide-bonded proteins are restricted to other compartments different from cytoplasmic space^28^, and bacteria are not an exception. In fact, in bacteria, there is no internal compartment. Only a few proteins use disulfide bonds as redox signalling systems such as OxyR and some reductases^29^. Only extracellular cysteines are oxidised to form disulfide bonds as part of their structural function, being the *fim* operon the most illustrative case^29^. All flagella protein are lacking disulfide bonds, and the amount of this amino acid is minimal^30^. In contrast, in cytoplasm most of the proteins are in reduced states thank the high level of reducing agents like glutathione that can reach values near 17 mM when *E. coli* is fed with glucose and growing in the exponential phase^31^. This study is a clear example of how the redox conditions of the environment constrain amino acid composition susceptibility. This fact could be acting concomitantly with the amino acid biosynthetic energetic cost^5^.

**Figure 3.**
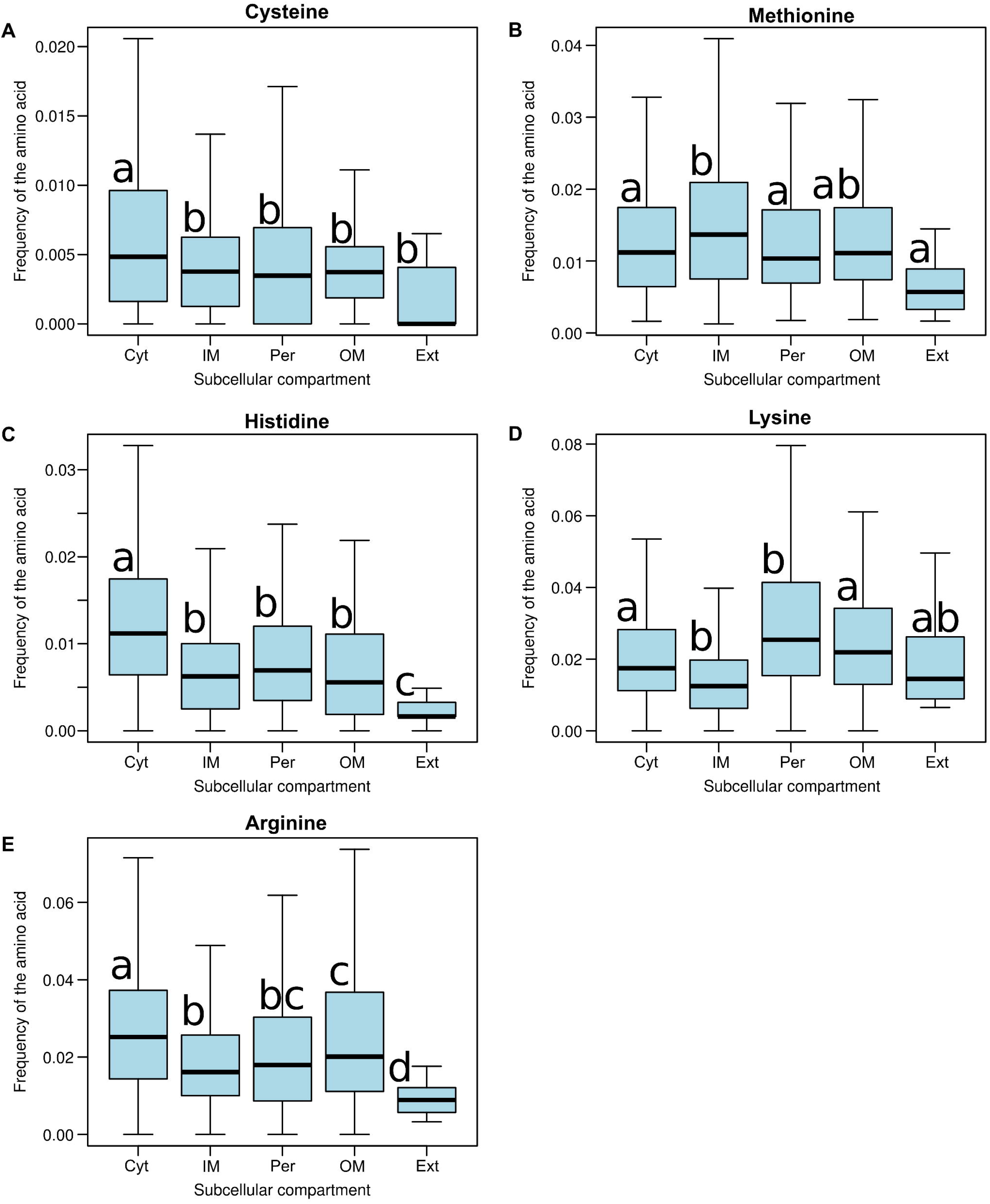
Frequency distribution of the most susceptible amino acids to oxidative damage: (A) cysteine, (B) methionine, (C) histidine, (D) lysine, and (E) arginine in all subcellular compartments (Cyt: cytoplasm, IM: inner membrane, Per: periplasm, OM: outer membrane, Ext: extracellular media). Different letters indicate significant differences, while the same one expresses no statistical differences (Nemenyi’s test). Note how cysteine, histidine and arginine drop their frequencies significantly from intracellular to the extracellular compartment.

### Methionine sulfoxidation

In contraposition to other amino acids, the oxidation of methionine is reversible being performed by the Msr family^32^. This reduction takes place in both the free form of amino acid and protein residues. We did not find any trend in the distribution of methionine among the subcellular compartments (GLM: slope = 9.385×10^-4^; *P* = 2.225×10^-2^; Spearman correlation: *r_s_* = 0.077; *P* = 6.151×10^-7^). There is a marginal difference in methionine composition between the cytoplasm and secretable proteins (Mann-Whitney *U* (cytoplasm – extracellular): *P* = 3.523×10^-2^) and no differences between the cytoplasm and periplasm and outer membrane (Mann-Whitney *U* (cytoplasm – periplasm): *P* = 6.045×10^-1^, Mann-Whitney *U* (cytoplasm – outer membrane): *P* = 2.812×10^-1^)) (Fig. 3B). Methionine residue oxidation can cause misfolding or rendering to dysfunctional proteins^33^. The methionine oxidation repair system, unique among amino acid oxidation repair damage, may contribute to the possibility of more extensive use of these amino acids in all compartments. Thus, methionine function is not easily replaceable, and the cells had to evolve the Msr family to continue using this amino acid in the presence of an oxidative environment. The methionine case provides good evidence that amino acid oxidative lesions could bias the amino acid frequency in proteins. We did not find differences in methionine oxidation, probably due to the activity of the repair system (Fig. 3B). The methionine oxidation repair highly converses among the tree of life^34^.

### Histidine oxidation

From the frequency in the protein sequence, we noted that the histidine is one of the third most scarce amino acids, after cysteine and tryptophan (Table S2). Histidine is used by proteins to coordinate metal atoms. We found that the cytoplasmic compartment had the highest level of histidine frequency while the extracellular one had the lowest level. In fact, there is a significant decrease in its content from intracellular to extracellular compartments (GLM: slope *=* - 5.583×10^-3^; *P* < 2.200×10^-16^; Spearman correlation: *r_s_* = −0.280; *P* < 2.200×10^-16^; Table S2, Fig. 3C). Even if the histidine frequency is compared between compartments, there is a significant difference between cytoplasmic proteins and the rest of compartments—from inner membrane to extracellular; Kruskal-Wallis Test: *P* < 2.200×10^-16^, Mann-Whitney *U* (cytoplasm – inner membrane): *P* = 3.125×10^-68^, Mann-Whitney *U* (cytoplasm – periplasm): *P* = 4.814×10^-16^, Mann-Whitney *U* (cytoplasm – outer membrane): *P* = 1.673×10^-9^, Mann-Whitney *U* (cytoplasm – extracellular): *P* = 5.955×10^-7^. For the oxidation products, histidine is the only amino acid that becomes into two different amino acids (asparagine and aspartic acid^35^). This phenomenon is similar to phenotypic mutations that eventually can undermine the information of the protein sequence. Probably, this consequence of the oxidative damage would occur proteome-wide under oxidative stress.

One of the roles of histidine inside the cell is the metal binding and coordination by some proteins^36^. Oxidative damage could lead to the evolution of siderophore biosynthetic pathways instead of histidine-based metal chelation systems as a strategy for metal assimilation such as iron, zinc, and manganese. Siderophores such as pyoverdine and catecholamine can protect the cell against UV- and antibiotic-derived ROS^37^. We could wander in the absence of oxygen, histidine-rich proteins would evolve as a preferential pathway replacing the function that siderophores play in microbial biology. Probably, oxygen undermined this possibility due to histidine sensitivity to ROS. Histidine-rich proteins were associated with bacterial habitat, being found mainly in rhizobia and pathogenic Gram-negative bacteria but not in obligate intracellular pathogens^38^. Some histidine-rich proteins such as ceruloplasmin and transferrin are involved in the chelation and transport of copper and iron, respectively^39,40^. One additional issue is that histidine oxidation could disrupt those signalling systems where histidine (de)phosphorylation play a fundamental role^41^.

### Aromatic amino acid oxidation

Aromatic amino acid residues are among the preferred targets for ROS attack^35^. However, there is even a slightly positive correlation between tyrosine, tryptophan and phenylalanine content and the spatial gradient (Tyr: GLM: Slope = 5.003×10^-3^; *P* < 2.200×10^-16^; Spearman correlation: *r_s_ =* 0.057; *P* = 2.560×10^-4^; Trp: GLM: Slope = 2.860×10^-3^; *P* = < 2.200×10^-16^; Spearman correlation: *r_S_* = 0.183; *P* < 2.200×10^-16^; Phe: GLM: Slope = 3.506×10^-3^; *P* = 4.152×10^-9^; Spearman correlation: *r_S_* = 0.129; *P* < 2.200×10^-16^). Tyrosine and tryptophan are the second and the fifth less abundant amino acid, respectively (Table S2). When the tyrosine frequency is compared between compartments, only there are significant differences between cytoplasmic proteins and the periplasm and the outer membrane—Mann-Whitney *U* (cytoplasm – periplasm): *P* = 3.599×10^-5^, Mann-Whitney *U* (cytoplasm – outer membrane): *P* = 4.138×10^-8^. However, the tryptophan frequency is unevenly distributed among compartments, probably due to the hydrophobic nature of this aromatic amino acid (Kruskal-Wallis test: *P* < 2.200×10^-16^, Mann-Whitney *U* (cytoplasm – inner membrane): *P* = 1.160×10^-43^, Mann-Whitney *U* (cytoplasm – periplasm): *P* = 6.422×10^-5^, Mann-Whitney *U* (cytoplasm – outer membrane): *P* = 6.783×10^-4^, Mann-Whitney *U* (cytoplasm – extracellular): *P* = 3.630×10^-2^). This uneven distribution was also found in the phenylalanine (Kruskal-Wallis test: *P* < 2.200×10^-16^, Mann-Whitney *U* (cytoplasm – inner membrane): *P* = 2.417×10^-84^, Mann-Whitney *U* (cytoplasm – periplasm): *P* = 2.195×10^-46^, Mann-Whitney *U* (cytoplasm – outer membrane): *P* = 6.482×10^-14^, Mann-Whitney *U* (cytoplasm – extracellular): *P* = 6.942×10^-9^). The aliphatic nature and the structural properties make them irreplaceable, explaining the lack of impact among compartments. These within-membrane amino acid over-representation is caused by the lack of polarity and compatibility in hydrophobic environments^42^.

### Peroxidation

Only valine, leucine, tryptophan and tyrosine are susceptible to peroxidation. While tryptophan and tyrosine are rare amino acids, leucine and valine are the most common and the fourth most abundant amino acids in *E. coli* (Table S2). In this subsection, we will focus more on leucine and valine as tryptophan and tyrosine were described in the previous paragraph. We detected that there were only significant differences in the frequency of leucine and valine between the cytoplasm and the inner membrane—Mann-Whitney *U* (Leu, cytoplasm – inner membrane): *P* = 1.835×10^-16^, Mann-Whitney *U* (Val, cytoplasm – inner membrane): *P* = 1.461×10^-5^. Peroxidation does not seem to have a fundamental impact on bias in the susceptible amino acid frequency. Maybe the nature of this reaction is so ubiquitous that natural selection partially solved evolving the scavenging systems, such as catalases and peroxidases, to limit general damage^43^.

### Carbonylation

Protein carbonylation is a type of protein oxidation promoted by ROS, and it refers to the formation of reactive ketones or aldehydes from the alcohol (-OH) groups in the side chain. Although all amino acids are susceptible to carbonylation if they are in the C-terminal of the protein, we focused on lysine, arginine, proline and threonine as they are more prone to be oxidised to carbonyl derivates^14, 22^. Nevertheless, the impact of carbonylation is not attributable only to oxidation to carbonyl groups, but may also be introduced into proteins by a mechanism that does not involve oxidation^14^. All amino acids showed differences in their frequencies between the cytoplasm and periplasm—Mann-Whitney *U* (Lys, cytoplasm – periplasm): *P* = 1.029×10^-14^, Mann-Whitney *U* (Arg, cytoplasm – periplasm): *P* = 1.284×10^-11^, Mann-Whitney *U* (Pro, cytoplasm – periplasm): *P* = 1.943×10^-2^, Mann-Whitney *U* (Thr, cytoplasm – periplasm): *P* = 1.451×10^-10^. Moreover, we detected significant differences in the frequency of lysine, arginine and proline between the cytoplasm and inner membrane—Mann-Whitney *U* (Lys, cytoplasm – inner membrane): *P* = 9.810×10^-38^, Mann-Whitney *U* (Arg, cytoplasm – inner membrane): *P* = 5.147×10^-40^, Mann-Whitney *U* (Pro, cytoplasm – inner membrane): *P* = 4.850×10^-4^—as well as differences in the frequency of lysine, arginine and threonine between the cytoplasm and outer membrane— Mann-Whitney *U* (Lys, cytoplasm – outer membrane): *P* = 1.574×10^-2^, Mann-Whitney *U* (Arg, cytoplasm – outer membrane): *P* = 4.537×10^-2^, Mann-Whitney *U* (Thr, cytoplasm – outer membrane): *P* = 4.774×10^-3^. Finally, differences in the frequency of arginine, proline and threonine were detected between the cytoplasm and extracellular proteins—Mann-Whitney *U* (Arg, cytoplasm – extracellular): *P* = 1.220×10^-5^, Mann-Whitney *U* (Pro, cytoplasm – extracellular): *P* = 1.275×10^-2^, Mann-Whitney *U* (Thr, cytoplasm – extracellular): *P* = 1.985×10^-3^ (Figs. 2 and 3D-E). These frequencies are coincident with expected decreased frequencies of amino acids prone to carbonylation from more reducing micro-environment (cytoplasm) to more oxidative ones such as extracellular compartments. Interestingly, we noticed that elevated frequencies of arginine and lysine were significantly higher in outer membrane proteins than in inner membrane ones (Figs. 3D-E). Arginine is more frequent in α-helix structural domains, which is also more abundant in outer membrane proteins while β-barrels are more common in inner membrane proteins^44^. A similar behaviour could be expected to happen with lysine due to its chemical similarity, although we did not find any report regarding this amino acid.

### Differences on the amino acid composition of inner transmembrane proteins

One special compartment is the inner membrane, where the same protein has amino acid residues exposed to the cytoplasmic reductive environment and periplasmic oxidative one at the same time. Thus, we paid particular attention to this situation and re-analysed the sequences within the same protein concerning amino acid frequency into the different protein segments (cytoplasm, transmembrane and periplasm). Amino acid composition of transmembrane proteins across regions reveals that there are significant differences in amino acid occurrence for all amino acids (Figure 4, Table S3). The disagreements among unevenly distributed amino acids across the three different locations within the same protein were significant. While alanine, cysteine, phenylalanine, isoleucine, leucine, valine, tryptophan and tyrosine are predominant in the transmembrane region, lysine and arginine are more common in the cytoplasmic region and aspartate, glycine, asparagine, proline, serine and threonine are more abundant in the periplasmic compartment (Figure 4, Table S3). This amino acid bias is mainly constrained to the protein secondary structure and function within the membrane^45,46^. Hence, there is not a clear pattern regarding amino acid distribution related to oxidation susceptibility.

**Figure 4.**
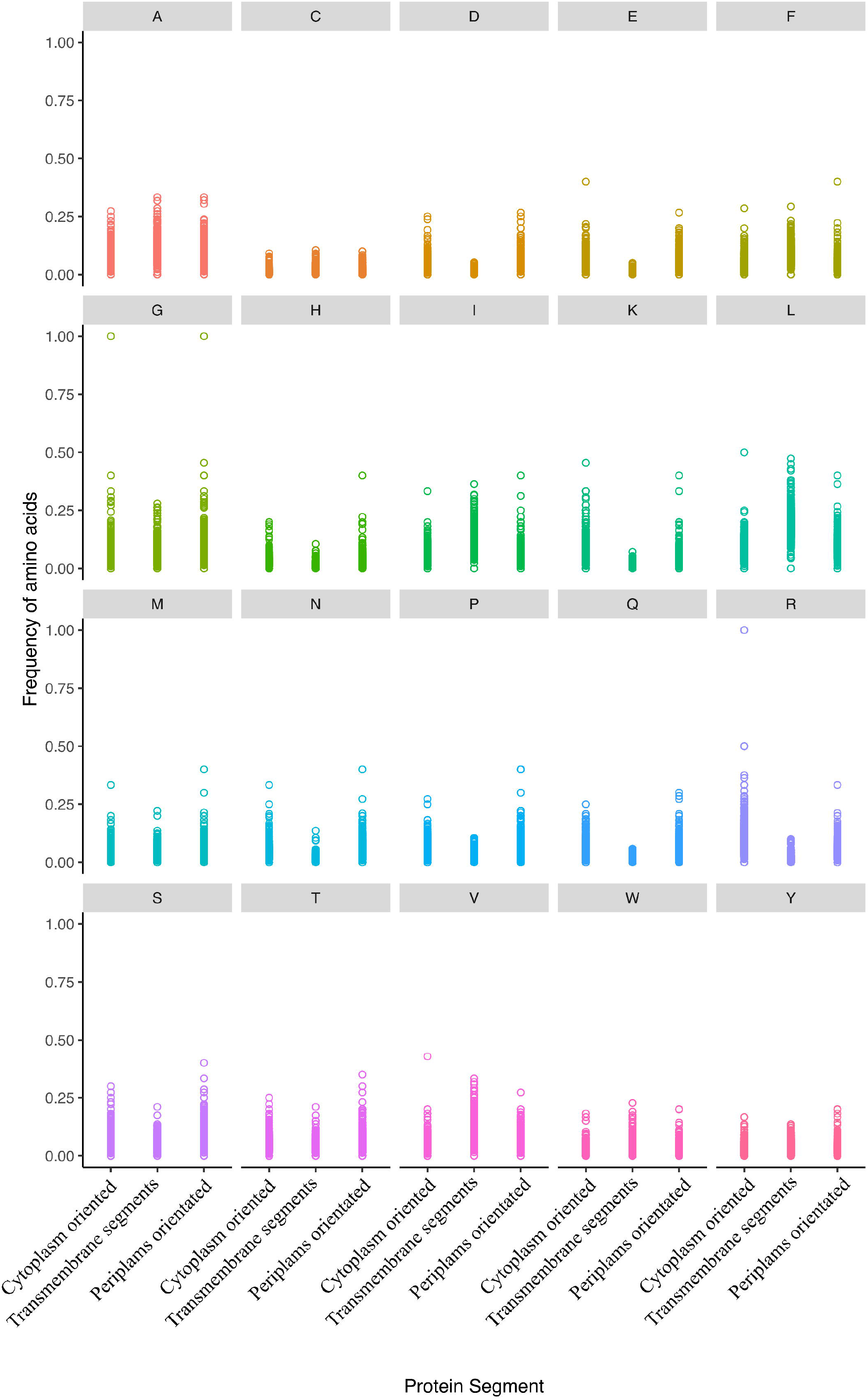
Distribution of the amino acid frequencies in inner protein fragments is segregated into three categories: cytoplasm-, transmembrane and periplasm-oriented. The plots represent the frequency of all amino acid across these categories. All frequencies are significantly different except for cysteine, histidine and lysine (Table S2).

### Conclusions

In the present study, we detected significant differences in amino acid frequencies in proteins across all cellular compartments for most of the amino acids. These data indicate an uneven distribution of amino acid that depends on the protein’s location within the cell in *E. coli*. This observation is consistent with a previous report which suggested that the amino acid distribution is related to the energetic cost^5^. Here, we propose that this amino acid bias in protein sequences could also be caused by individual amino acid residue susceptibility to different oxidative lesions. We found that primary susceptible amino acids are overrepresented in the cytoplasm and decreased among the different subcellular compartments, being correlated with the redox gradient from reductive to oxidative microenvironments. The uneven distribution of amino acids across cellular compartments depends on multiple components, such as structural constraints, catalytic amino acids, subcellular compartment polarity, and protein information.

## Materials and methods

### Data mining

The whole proteome of *Escherichia coli* K-12 MG1655, consisting of 4143 proteins, was downloaded from EcoGene 3.0^47^ on 26/03/2015 in FASTA format. Every protein was analysed to count the number of each amino acid using a custom Python script. A tabular-separated values (tsv) file where every file is a single protein, and every column represents the absolute frequency of amino acid in such protein was obtained. The susceptibility of the amino acids to the different oxidative stresses was based according to a previous classification^22^.

### Distribution of the amino acid composition per subcellular compartment

As protein size distribution did not fit a normal distribution^48^, values were presented as the frequency of amino acid normalised by the median size of proteins in every cellular compartment. We consider the classical protein definition and only polypeptides larger than 50 amino acid^49^ were considered for all analysis. To test if the content of every amino acid among proteins between the different cellular compartments (i.e. cytoplasm, inner membrane, periplasm, outer membrane and secreted) is uniformly distributed, Kruskal-Wallis test was performed. If the differences are significant, multiple comparisons using the Nemenyi’s test^50^ were made. False discovery rate^51^ was used to correct *P*-values of *posthoc* comparisons in all these tests. Bonferroni corrected Mann-Whitney *U* was performed for pair comparisons as indicated across the text. Finally, to describe the spatial gradient of amino acid oxidative lesions, generalised linear models (GLM) per amino acid and subcellular compartment and Spearman correlation were done. All these analyses were made in R 3.2.1^52^ using the *HH^53^* and *PMCMR^54^* packages.

### Distribution of the amino acid composition in transmembrane proteins

The prediction of transmembrane regions was made using Phobius 1.01^55^ and TMHMM 2.0^56^. All signal peptides were checked with Signal-P 4.0^57^ and removed in these analyses. All analyses related to the distribution of the amino acids in the different regions were performed as described previously.

## Acknowledgements

We thank to Arpita Nath, Dr. Dan Roizman and Dr. Flor I. Arias-Sánchez (from Freie Universität Berlin) for valuable comments on the manuscript. We also acknowledge Prof. Dr. Jens Rolff for …

---

**Table S1. Original data set of *Escherichia coli* proteins and its amino acid composition modified from EcoGene v. 3.0^47^.** Polypeptides smaller than 50 amino acids and pseudogene which original location are unknown were excluded from further analyses.

**Table S2. Amino acid composition and distribution of the *E. coli* proteome across subcellular compartments.** Numbers in the first five fields indicate the median of the normalised frequencies of every amino acid per the median protein length in every compartment. Different letter(s) state(s) significant differences (*P* < 0.05) between compartments according to Nemenyi’s test and after applying false discovery rate. Spearman *r_s_* and *P*-values indicate the correlation between the amino acid frequency and the subcellular compartments and its significance, respectively. Finally, the slope, the coefficient of determination (*R*^2^) and the GLM *P*-value reflects the trend of the amino acid distribution across subcellular compartments, the adjust degree to a generalised linear model and its significance. All *P* values shown in bold are significant after Bonferroni correction.

**Table S3. Amino acid composition of the different sections of all *E. coli* transmembrane proteins.** Numbers in every field indicate the median of the normalised frequencies of every amino acid per the median protein length in every compartment. Different letter(s) show(s) significant differences (*P* < 0.05) between compartments according to Nemenyi’s test and after applying false discovery rate. Spearman *r_s_* and *P* values indicate the correlation between the amino acid frequency and the subcellular compartments and its significance, respectively. Finally, the slope, the coefficient of determination (*R*^2^) and the GLM *P*-value reflects the trend of the amino acid distribution across subcellular compartments, the adjust degree to a generalised linear model and its significance. All *P* values shown in bold are significant after Bonferroni correction.

